# VStrains: De Novo Reconstruction of Viral Strains via Iterative Path Extraction From Assembly Graphs

**DOI:** 10.1101/2022.10.21.513181

**Authors:** Runpeng Luo, Yu Lin

## Abstract

With the high mutation rate in viruses, a mixture of closely related viral strains (called viral quasispecies) often co-infect an individual host. Reconstructing individual strains from viral quasispecies is a key step to characterizing the viral population, revealing strain-level genetic variability, and providing insights into biomedical and clinical studies. Reference-based approaches of reconstructing viral strains suffer from the lack of high-quality references due to high mutation rates and biased variant calling introduced by a selected reference. De novo methods require no references but face challenges due to errors in reads, the high similarity of quasispecies, and uneven abundance of strains.

In this paper, we propose VStrains, a de novo approach for reconstructing strains from viral quasispecies. VStrains incorporates contigs, paired-end reads, and coverage information to iteratively extract the strain-specific paths from assembly graphs. We benchmark VStrains against multiple state-of-the-art de novo and reference-based approaches on both simulated and real datasets. Experimental results demonstrate that VStrains achieves the best overall performance on both simulated and real datasets under a comprehensive set of metrics such as genome fraction, duplication ratio, NGA50, error rate, *etc*.

**Availability:** VStrains is freely available at https://github.com/MetaGenTools/VStrains.

## 1 Introduction

Viruses are the most abundant biological entities on Earth and have high mutation rates, up to a million times higher than their hosts [11,26]. Variations in viral genetic sequences lead to the emergence of new viral strains during evolution and are also known to be associated with many diseases [31]. One challenge in viral studies is to analyze a mixture of closely related viral strains, referred to as *viral quasispecies*. The problem of inferring individual viral strains from sequencing data of viral quasispecies is called *strain-aware assembly* or *viral haplotype reconstruction*. Viral haplotype reconstruction in individual patients provides a signature of genetic variability and thus informs us about disease susceptibilities and evolutionary patterns of distinct viral strains [10]. Sequencing data of viral quasispecies from next-generation sequencing (NGS) techniques are short and have a low error rate [33] (≤0.5%) while that from third-generation sequencing (TGS) techniques are long and have a high error rate [9] (1.6–2.7% for deletions, 1.2–2.2% for mismatches and 1.1–2.4% for insertions). Due to the low pairwise strain divergence in viral quasispecies, it is challenging to distinguish the sequencing error in TGS data and highly similar viral strains. Therefore, various approaches have been proposed to infer individual viral strains from NGS data and can mainly be classified into two categories [15], reference-based and de novo (or reference-free). Reference-based approaches (such as PredictHaplo [30] and NeurHap [36]) rely on the alignment between reads and references and thus suffer from the lack of high-quality references due to high mutation rates [8], and biased variant calling introduced by a selected reference [3,34]. De novo approaches directly assemble viral strains from sequencing reads without references and have the potential to identify novel viral strains and provide deep insights for viral genetic novelty [31].

While de novo (meta)-genomic assemblers such as SPAdes-series [1,5,7,24,28] could be applied to assemble individual strains, they tend to produce fragmented contigs rather than complete viral strains, or collapsed contigs ignoring differences between strains, as they are not specifically designed to distinguish closely related viral strains. Specialized de novo assemblers such as SAVAGE [3], PEHaplo [8], viaDBG [12] and Haploflow [13] directly assemble reads into strains from viral quasispecies and have achieved promising results. More recently, VG-Flow [4] was proposed to extend pre-assembled contigs (produced by the specialized assembler SAVAGE [3]) into full-length viral strains using flow variation graphs and significantly outperformed other approaches on recovering viral strains from viral quasispecies. While VG-Flow [4] guarantees its runtime to be polynomial in the genome size, all the recovered viral strains must be selected from a set of candidate paths inferred by greedy path extraction strategies and thus some viral strains not covered by the candidate set are infeasible to be reconstructed.

Here we propose VStrains, a de novo approach for reconstructing strains from viral quasispecies. VStrains employs SPAdes [5] to build the assembly graph from paired-end reads and incorporates contigs and coverage information to iteratively extract distinct paths as reconstructed strains. We benchmark VStrains against multiple state-of-the-art de novo and reference-based approaches on both simulated and real datasets. Experimental results demonstrate that VStrains achieves the best overall performance on both simulated and real datasets under a comprehensive set of metrics such as genome fraction, duplication ratio, NGA50, error rate, etc. In particular, in more challenging real datasets, VStrains achieves remarkable improvements in recovering viral strains compare to other methods.

## 2 Methods

### 2.1 Preliminary

An assembly graph generated by SPAdes is a directed graph *G* = (*V, E*). Each vertex *v* ∈ *V* represents a double-stranded DNA segment where its forward and reverse strands are denoted by *seq*(*v*^+^) and *seq*(*v*^*−*^), respectively. A distinctive feature of assembly graphs built by SPAdes (with iterative *k*-mer sizes up to *k*_*max*_) is that each (*k*_*max*_+1)-mer appears in at most one vertex and thus can be used to uniquely identify the corresponding vertex in the assembly graph. Two vertices *u* and *v* can be connected by an edge 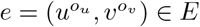, where *o*_*u*_, *o*_*v*_ ∈ {+,−} denote the strandedness of *u* and *v*, respectively. Note that the suffix of 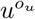 overlaps *k*_*max*_ positions with the prefix of 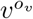 under the de Bruijn graph model [29] behind SPAdes. Moreover, the assembly graphs built by SPAdes also contain contigs information, *i*.*e*., a contig in *G*(*V, E*) is defined as a path of vertices together with their strandedness information. The *coverage* of a vertex *v* is estimated by the number of reads containing the DNA segment corresponding to *v* and denoted by *cov*(*v*). The coverage of a contig *c* is estimated by the average coverage along all the vertices in *c* and denoted by *cov*(*c*).

### 2.2 Algorithm Overview

VStrains takes paired-end reads from viral quasispecies as the input and aims to recover individual viral strains. During pre-processing, VStrains first employs SPAdes to construct an assembly graph and contigs from paired-end reads, then canonizes the strandedness of vertices and edges, and further complements the assembly graph with additional linkage information from paired-end reads. After pre-processing, VStrains makes use of the contigs, paired-end links, and coverage information to perform branch splitting and non-branching path contraction to disentangle the assembly graph. Finally, VStrains outputs strain-specific sequences from the assembly graph via iterative contig-based path extraction. Refer to Fig. 1 for an overview of our algorithm. Details of each step are explained in the following sections.

**Fig. 1:**
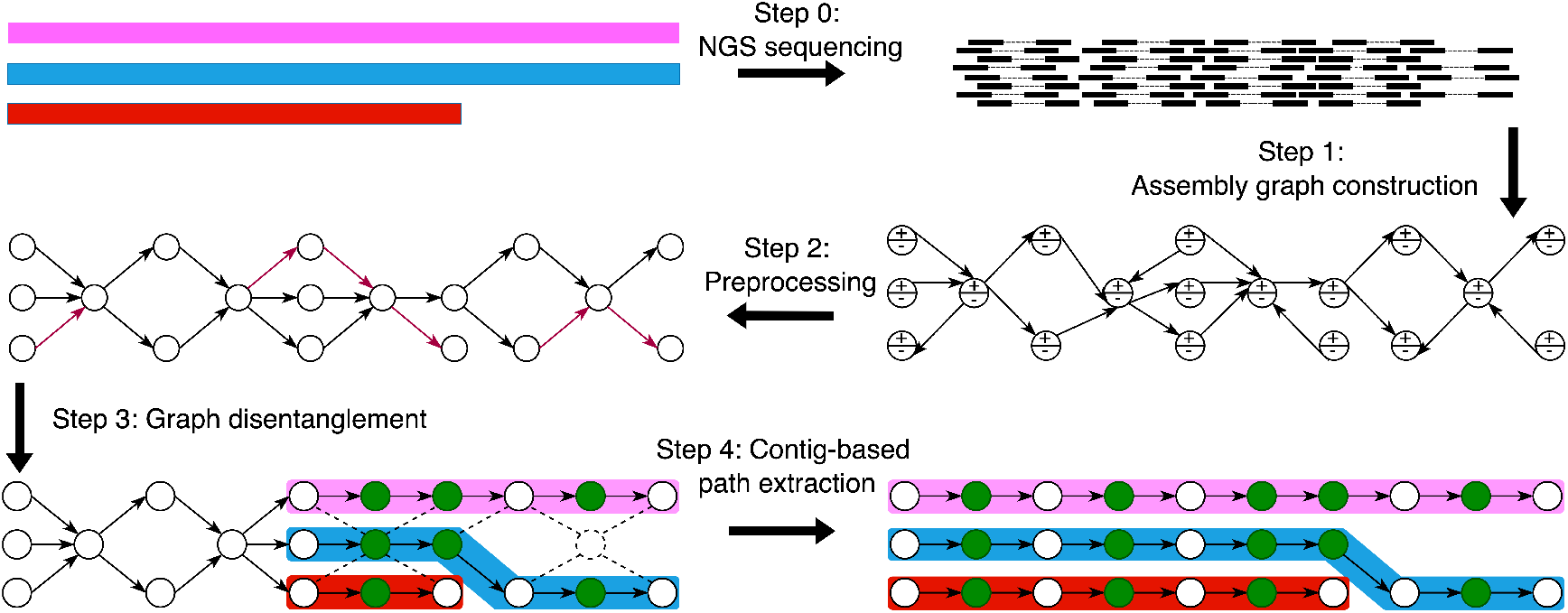
The framework of VStrains

### 2.3 Preprocessing

#### 2.3.1 Canonize Strandedness

Recall that each vertex in the assembly graph produced by SPAdes represents both the forward and reverse strands of a DNA segment, and thus contigs reported by SPAdes may refer to two different strands of the same DNA segment. Therefore, there is no obvious correspondence between a viral strain and a directed path in the assembly graph. To unravel this correspondence, VStrains performs the strandedness canonization to choose the strandedness *o*_*v*_ ∈ {+, −} for each vertex *v* ∈ *V* where all adjacent edges only use 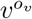. VStrains first chooses an arbitrary vertex *s* ∈ *V* as the *starting vertex* and fixes its strandedness to *o*_*s*_. Let 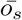 denotes the opposite strandedness of *o*_*s*_, i.e., 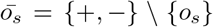. VStrains flips its adjacent edges if necessary to ensure these edges only use 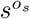, e.g., 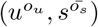 and 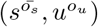 will be flipped into 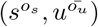 and 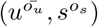, respectively. VStrains iteratively performs the above step until all the vertices have a fixed strandedness or one vertex has to use both strandedness. VStrains resolves the latter case by splitting this vertex into a pair of vertices, representing its forward and reverse strands, respectively. As a result, *seq* 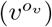can be simplified to *seq*(*v*) where *o*_*v*_ is the chosen strandedness of *v*, and 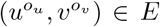 can be simplified to (*u, v*), where *o*_*u*_ and *o*_*v*_ are the chosen strandedness of *u* and *v*, respectively. The vertices without any in-coming edges are defined as *source* vertices, whereas the vertices without any out-going edges are defined as *sink* vertices.

#### 2.3.2 Inferring Paired-end Links

While SPAdes uses paired-end reads to construct contigs as paths in the assembly graph, VStrains uses unique *k*-mers of vertices in the assembly graph to establish mappings between paired-end reads and pairs of vertices and thus infers paired-end links.

VStrains uses minimap2 [20] to find exact matches of (*k*_*max*_+1)-mers between pairs of vertices in the assembly graph produced by SPAdes (with iterative *k*-mer sizes up to *k*_*max*_) and paired-end reads. Assume *u* and *v* are a pair of vertices in the assembly graph. A *PE* link is added between *u* and *v* if a paired-end read contains at least one (*k*_*max*_+1)-mer in *u* and at least one (*k*_*max*_+1)-mer in *v*. Note that the (*k*_*max*_+1)-mer can uniquely identify the corresponding vertex and thus makes it extremely unlikely to produce false-positive *PE* links unless errors in reads coincide with rare variations between strains. Since (*k*_*max*_+1)-mer is usually smaller than the read length, even erroneous paired-end reads may infer paired-end links as long as they still contain error-free (*k*_*max*_+1)-mers.

Note that paired-end reads are commonly used to infer paired-end links between vertices in the assembly graph. For example, overlap-graph-based assemblers, such as SAVAGE [3] and PEHaplo [8], use pairwise alignments between paired-end reads to build overlap graphs [27], and thus face challenges to choose appropriate parameters (*e*.*g*., the overlap length cutoff along with allowed mismatches) to distinguish false-positive links between perfect reads from different strains and true positive links between erroneous reads from the same strain in overlap graphs. De Bruijn graph-based assemblers, such as SPAdes and viaDBG, split paired-end reads into *k*-bimers [5] or bilabel [23], pairs of *k*-mers of exact (or near exact) distances, one from the forward read and the other from the reverse read. Although this split strategy helps adjust *k*bimers/bilabel distances [5,23] and efficiently construct de Bruijn graphs, it does not make full use of all available *k*-mer pairs (with varying distances) between forward and reverse reads to further simplify their assembly graphs. To overcome this limitation, VStrains uses all available *k*-mer pairs without any distance constraints between forward and reverse reads to create *PE* links between vertices in the assembly graph, which unravels its potential to further simplify the assembly graph produced by SPAdes.

### 2.4 Graph Disentanglement

After pre-processing, paired-end link information has been incorporated, and viral strains are expected to correspond to directed paths in the assembly graph. However, strains may share vertices and edges, and thus result in an entangled assembly graph. In this section, the graph disentanglement iteratively splits branching vertices and contracts non-branching paths as follows.

#### 2.4.1 Branching vertex splitting

A vertex *v* ∈ *V* is called a *branching vertex* if either the in-degree or out-degree of *v* is greater than 1. A branching vertex is *non-trivial* if both its in-degree and out-degree are greater than 1, and *trivial* otherwise. For example, vertex L is a trivial branching vertex while vertices D, G, K, and P are non-trivial branching vertices in Fig. 2(a).

**Fig. 2:**
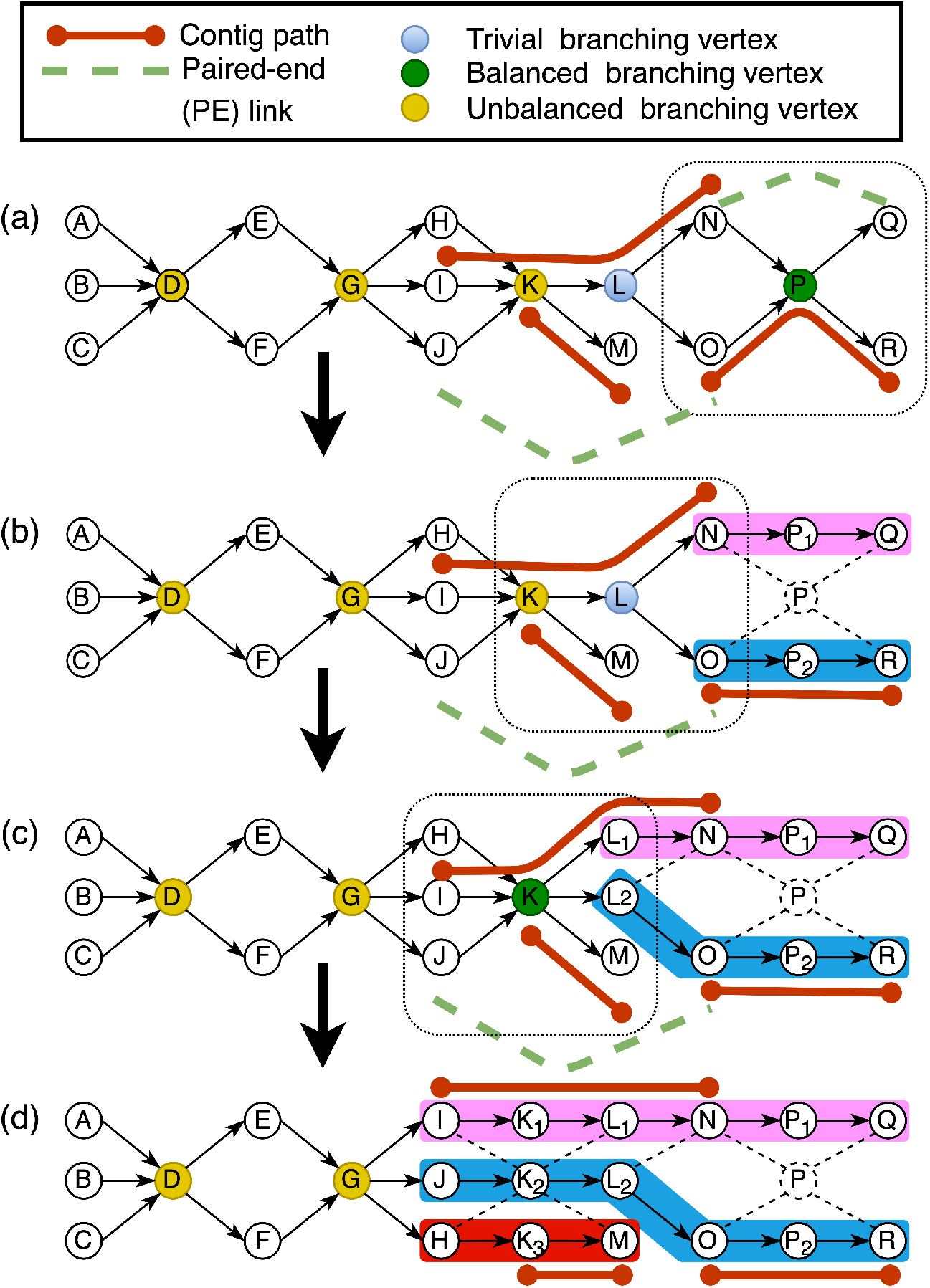
Graph disentanglement of VStrains. (a) P is a balanced branching vertex. (b) P is split into P_1_ and P_2_ in a balanced split corresponding to O↔R(contig path) and N↔Q (*PE* link). (c) A trivial branching vertex L in (b) is split into L_1_ and L_2_ in a trivial split, and thus previously unbalanced branching vertex K becomes a balanced branching vertex. (d) K is split into K_1_, K_2_ and K_3_ in a balanced split, corresponding to three one-to-one mappings, I↔L_1_NP_1_Q (contig path), J↔L_2_OP_2_R (*PE* link), and H↔M (coverage compatible pair). D and G are not split during graph disentanglement.

Without loss of generality, assume a trivial branching vertex *v* has multiple in-coming edges {(*u*_*i*_, *v*) ∈ *E* |*i* = 1, …, *n*}. In a trivial split, vertex *v* will be replaced by vertices {*v*_1_, …, *v*_*n*_}, and each in-coming edge (*u*_*i*_, *v*) will be replaced by (*u*_*i*_, *v*_*i*_), respectively. The coverage of *v*_*i*_ and the capacity of (*u*_*i*_, *v*_*i*_) are set to the capacity of (*u*_*i*_, *v*). If *v* has an out-going edge (*v, w*), (*v, w*) will be replaced by edges {(*v*_*i*_, *w*)|*i* = 1, …, *n*} where the capacity of (*v*_*i*_, *w*) is set to the coverage of *v*_*i*_.

A non-trivial branching vertex *v* is called *balanced* if *v* has the same number of in-coming edges {(*u*_*i*_, *v*) ∈ *E* |*i* = 1, …, *n*} and out-going edges {(*v, w*_*j*_) ∈ *E* |*j* = 1, …, *n*}. For example, vertex P is a balanced branching vertex in Fig. 2(a). Let *U* = {*u*_*i*_|*i* = 1, …, *n*} and *W* = {*w*_*j*_|*j* = 1, …, *n*}. In a balanced split, the balanced branching vertex *v* will be replaced by vertices {*v*_1_, …, *v*_*n*_}, and an in-coming edge (*u*_*i*_, *v*) and out-going edge (*v, w*_*i*_) will be replaced by (*u*_*i*_, *v*_*i*_) and (*v*_*i*_, *w*_*i*_) if *u*_*i*_ and *w*_*i*_ are both contained in at least one contig or connected by a *PE* link. The above balanced split of *v* corresponds to a bijection between *U* and *W*, and most of these one-to-one mappings can be perfectly inferred by contigs and *PE* links between *U* and *W*. For example, a balanced split of P from Fig. 2(a) to (b) corresponds to two one-to-one mappings, O↔R and N↔Q, inferred by the contig and *PE* link information, respectively. In case *U* and *W* form a partial bijection using contigs and *PE* links information, VStrains further uses coverage information and aims to find more one-to-one mappings. For the *u*_*i*_ ∈ *U* and *w*_*j*_ ∈ *W* not in the current partial bijection, an one-to-one mapping is established between *u*_*i*_ and *w*_*j*_ if and only if *u*_*i*_ and *w*_*j*_ form a *coverage compatible pair, i*.*e*., 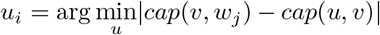 and 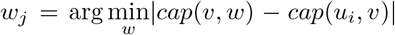. For example in Fig. 2(c) to (d), vertices H and M form a coverage compatible pair, together with two other one-to-one mappings I↔L_1_NP_1_Q (from contig path) and J ↔ L_2_OP_2_R (from *PE* link), lead to a balanced split on the balanced branching vertex K in Fig. 2(d).

Note that not all non-trivial branching vertices are balanced (*e*.*g*., vertex K is an unbalanced branching vertex in Fig. 2(a)). For such unbalanced vertex *v*, VStrains performs a trivial split on its adjacent trivial branching vertices and aims to convert *v* into a balanced vertex. For example, after performing a trivial split on vertex L in Fig. 2(b) to (c), vertex K now becomes a balanced branching vertex in Fig. 2(c) and becomes a candidate for a balanced split. VStrains performs a balanced split on *v* if the above bijection can be established by contigs, *PE* links, and coverage information.

#### 2.4.2 Non-branching path contraction

The above branch split operation in the assembly graph creates non-branching paths, i.e., path *p* = (*v*_1_, *v*_2_, …, *v*_*n*_), *v*_*i*_ ∈ *V* ∀*i* = 1, …, *n*, where the in-degree of *v*_2_, …*v*_*n*_ and the out-degree of *v*_1_, …*v*_*n−*1_ are all 1. Following the similar idea of graph simplification in SPAdes, VStrains contracts all the non-branching paths. For example, the above non-branching path *p* is contracted into one vertex *v*_*p*_, and each in-coming edge (*u, v*_1_) of *v*_1_ is replaced by (*u, v*_*p*_) and each out-going edge (*v*_*n*_, *w*) is replaced by (*v*_*p*_, *w*) with the same capacity, the coverage of *v*_*p*_ is set to be the average coverage of *v*_1_, …, *v*_*n*_. Moreover, *v*_*p*_ inherits all the *PE* links of vertices in non-branching path *p*. For example in Fig. 2(d), three non-branching paths in three different colors on the right are contracted, respectively.

### 2.5 Contig-based Path Extraction

While VStrains effectively disentangles the assembly graph through branching vertex splitting and non-branching path contraction in the above step, there still exist branching vertices (*e*.*g*., D and G in Fig. 2(d)) which may introduce ambiguity in distinguishing full paths that correspond to individual viral strains.

Recall SPAdes outputs contigs as paths of vertices on the assembly graph, which are usually sub-paths of viral strain induced paths on the assembly graph. The remaining problem is to extend these contigs (sub-paths) into corresponding viral strains (full paths) on the assembly graph. Note that a full path on the assembly graph starts from a source vertex and ends at a sink vertex or is a cyclic path (*i*.*e*., its first and the last vertex coincide). Ideally, the strain-specific (not shared by multiple viral strains) contigs should be extended and extracted first. The longer a contig is, the more likely that this contig is strain-specific.

Therefore, VStrains iteratively selects the longest contig and extends its corresponding subpath on both ends. Without loss of generality, consider the right extension of the current contig *C* = (*v*_1_, *v*_2_, …, *v*_*n*_). If *v*_*n*_ has only one out-going edge (*v*_*n*_, *v*_*n*+1_), the current contig will be extended into (*v*_1_, …, *v*_*n*_, *v*_*n*+1_). If *v*_*n*_ has multiple out-going edges 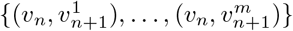, we follow the same strategy in Section 2.4.1 to look for one-to-one mapping between *v*_*n−*1_ and 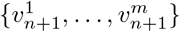 using contigs, paired-end reads and coverage information. If *v*_*n−*1_ forms the one-to-one mapping with 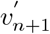, the current contig will be extended into 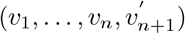, otherwise, the extension to the right terminates. If *v*_*n*_ is a sink vertex (without any out-going edges), the extension to the right terminates. If *v*_*n*_ is a visited vertex during extension on the other end, the extension to the right terminates and a cyclic path is obtained by combining both left and right extensions.

If the currently selected contig can be extended into a full path, VStrains will include this extended path as one of the output strains. Otherwise, VStrains will contract this extended path as a single vertex, and wait for confident extension in the future step. VStrains will also update the coverage information of the assembly graph. More specifically, VStrains estimates the path coverage as the median coverage of all the non-branching vertices along the path, and reduces the path coverage from all the traversed vertices. For example, VStrains extends the contig H→K_3_→M into the full path C→D_1_→ … →K_3_→M (using coverage compatible pairs F↔H and C↔F described in Section 2.4.1) in Fig. 3(b).

**Fig. 3:**
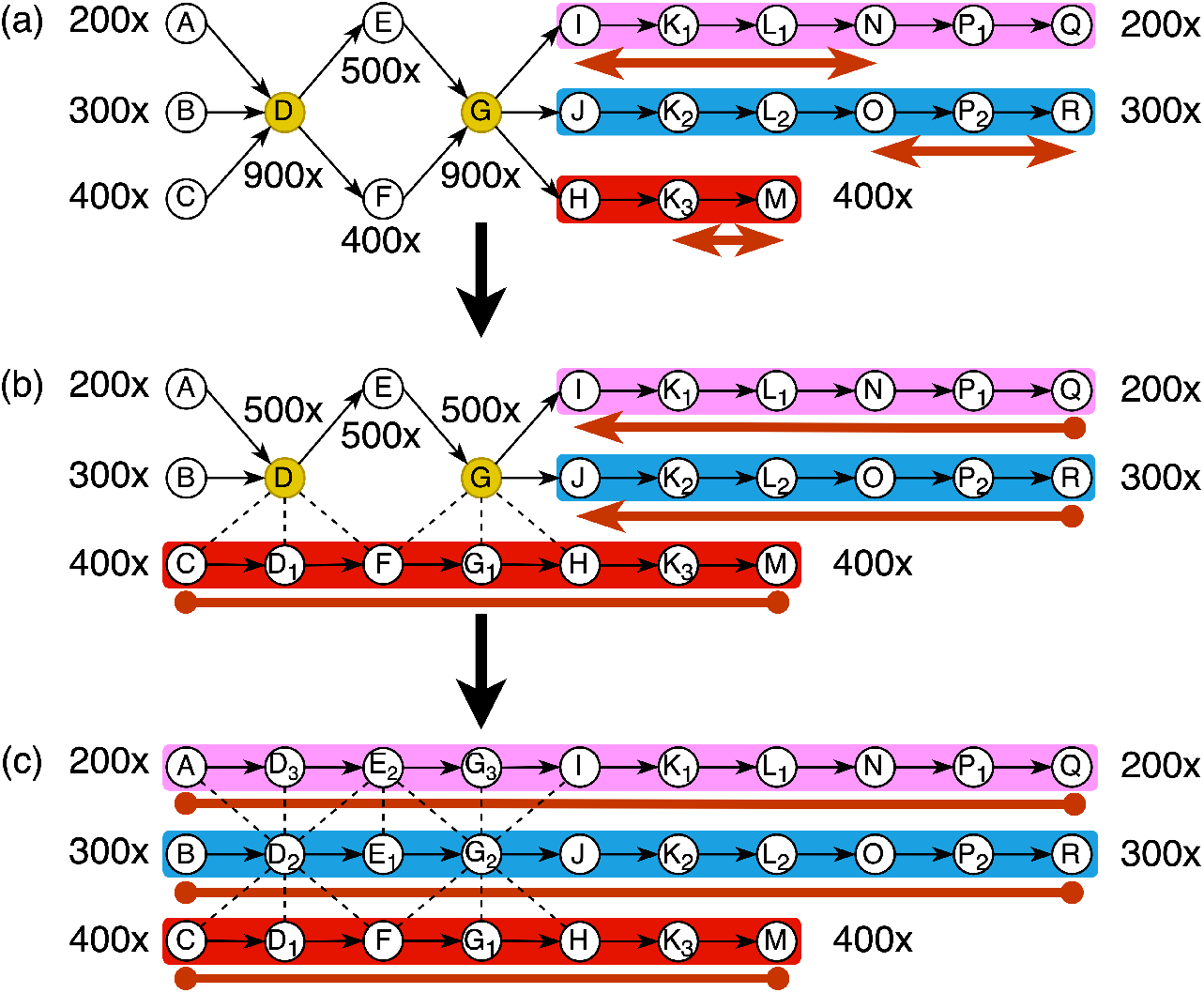
The contig-based path extraction of VStrains. (a) Assembly graph after graph disentanglement with three contigs: I→K_1_→L_1_→N, O→P_2_→R, K_3_→M. (b) The longest contig I→K_1_ L_1_→N is firstly extended to I→K_1_… P_1_→Q (terminated at I due to the lack of coverage compatible pair with respect to 200x). Afterwards, the second longest contig O→P_2_→R is extended to J→K_2_→ … →P_2_→R (terminated at J due to the lack of coverage compatible pair with respect to 300x). At last, the shortest contig K_3_→M is extended to C→D_1_→ … →K_3_→M (thanks to the coverage compatible pairs with respect to 400x) (c) Contigs J→ … →R and I→ … →Q are extended to full paths (thanks to the path extraction and coverage/topology update in (b)).

By iteratively extending and extracting the contig from the assembly graph, VStrains obtains a set of distinct paths as the final viral strains. This iterative strategy contracts/extracts the most confident sub-paths/full-paths first, and updates topology and coverage information of the assembly graph on the fly, which in turn reveals more strain-specific paths in the updated graph and facilitates the subsequent path extractions. For example, after extracting the full path C→D_1_→ … →K_3_→M in Fig. 3(b), the (sub)-paths I→K_1_→ … →P_1_→Q and J→K_2_→ … →P_2_→R are able to further extend to the left (using coverage compatible pairs A↔I and B↔J) into two full paths A→D_3_→ … →P_1_→Q and B→D_2_→ … →P_2_→R.

Note that, unlike other greedy path finding strategies [4,8,13] deriving a set of candidate paths based on the original flow-variation graph, VStrain iteratively extracts the most confident path and updates the assembly graph on the fly, which results in more accurate reconstruction of viral strains. For example, VG-Flow employs three greedy strategies (maximum capacity, minimum capacity, shortest paths) to derive the set of candidate paths, from which the final output strains will be selected. However, such greedy strategies are directly applied on the original flow-variation graph, making it likely to find erroneous paths (e.g., the maximum-capacity path C→D→E→G→H→K_3_→M and the minimum-capacity path A→D→F→G→I→ … →P_1_→Q in Fig. 3(a) are both erroneous paths).

## 3 Experimental Setup

### 3.1 Experimental Datasets

#### 3.1.1 Simulated Datasets

To evaluate the performance and scalability of VStrains, we used three simulated viral quasispecies datasets from [3] consisting of 6 Poliovirus, 10 hepatitis C virus (HCV), and 15 Zika virus (ZIKV) mixed strains, respectively. These datasets were commonly used to benchmark strain-aware viral assemblers and simulated from known reference genomes using SimSeq [6] with the default error profile.

#### 3.1.2 Real Datasets

Two real datasets with different coverages are obtained from the NCBI database. The first 5-HIV-labmix (20,000x) dataset [14] is a lab mix of 5 known real human immunodeficiency virus (HIV) strains (NCBI accession number *SRR961514*). The second 2-SARS-COV-2 (4,000x) dataset [37] is a mixture of 2 real severe acute respiratory syndrome coronavirus 2 (SARS-COV-2) strains (BA.1 and B.1.1). Two independently assembled strains, BA.1 (NCBI accession number *SRR18009684*) and B.1.1 (NCBI accession number *SRR18009686*), are used as the ground-truth for evaluation. Table 1 presents a summary of the simulated and real datasets.

**Table 1:**
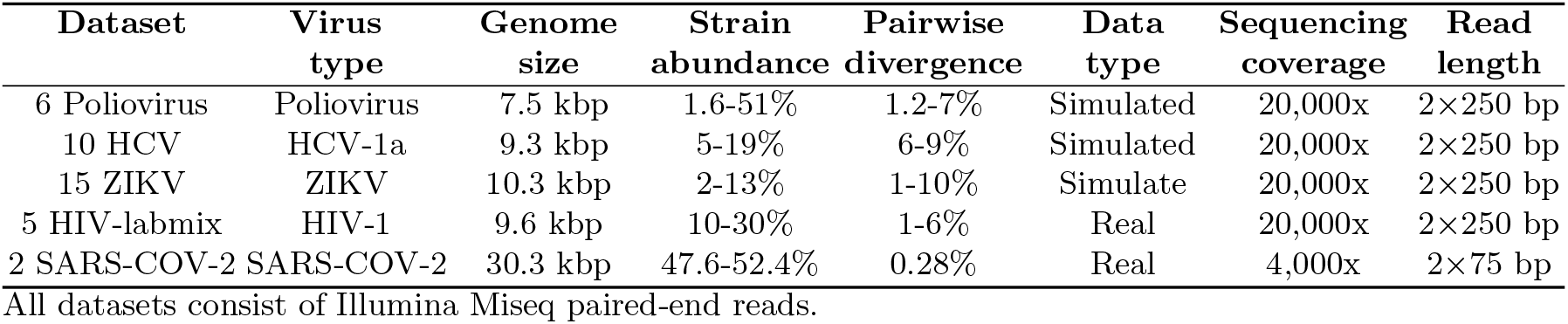
Quasispecies characteristics of the simulated and real benchmarking datasets.

### 3.2 Baselines and Evaluation Metrics

VStrains was benchmarked against the state-of-the-art methods on assembling viral strains including SPAdes [5], SAVAGE [3], VG-Flow [4], PEHaplo [8], viaDBG [12], Haploflow [13], and PredictHaplo [30]. Three recent reference-based machine-learning approaches GAEseq [17], CAECseq [16] and NeurHap [36] were not included for comparison because all of them have only been applied to a gene segment of viral strains in their experiments and failed to handle the above datasets on whole viral strains (*i*.*e*., memory usage exceeding 300GB). Note that PredictHaplo [30] is a reference-based approach and needs an accurate reference as the input. Therefore, we randomly select a ground-truth viral strain and provide it to PredictHaplo [30] for each dataset. All above tools under evaluation use default settings unless specified otherwise. Refer to Section S3 in the Supplementary Materials for a detailed description of baselines.

Similar to previous studies, we employ MetaQUAST [25] to evaluate all assembly results of viral strains. As viral quasispecies contains highly similar strains, we run MetaQUAST with the “--unique-mapping” option to minimize ambiguous false positive mapping. For each assembly, we report the genome fraction, duplication ratio, NGA50, error rate, and number of contigs. Genome fraction is defined as the total number of aligned bases in the reference, divided by the genome size. A base in the reference genome is counted as aligned if there is at least one contig with at least one alignment to this base. Duplication ratio is defined as the total number of aligned bases in the assembly, divided by the total number of aligned bases in the reference. NG50 is the contig length such that using longer or equal length contigs produce half of the bases of the reference genome, whereas NGA50 counts the lengths of aligned blocks instead of contig lengths, such that the contig is broken into smaller pieces when it has a misassembly with respect to the reference genome. Error rate is defined as the sum of mismatch rate, indel rate, and N’s rate, which reflects the number of errors with respect to the reference genome size.

## 4 Experimental Results

In this section, we show the performance of VStrains and other baselines on both simulated and real datasets. In consistent with previous observations [4,12], we found that SPAdes in general outperforms metaSPAdes [28] and other specialized versions (refer to Section S1 in the Supplementary Materials for detailed comparison among SPAdes-series assemblers) and thus is employed to build assembly graphs for VStrains.

### 4.1 Performance on Simulated Datasets

Table 2 summarizes the performance of de novo and reference-based approaches on reconstructing viral strains in three simulated datasets. Note that specialized strain-aware assemblers such as PEHaplo, viaDBG, and SAVAGE typically outperform the general-purpose assembler SPAdes, especially in genome fraction. One possible reason is that SPAdes is not designed to distinguish highly similar viral strains. When two or more strains share long and identical sequences, SPAdes may result in fragmented assemblies (*i*.*e*., low NGA50) and keep only one copy of such shared sequences (*i*.*e*., low genome fraction). VG-Flow uses assembled contigs from SAVAGE (VG-Flow+SAVAGE) and SPAdes (VG-Flow+SPAdes) to build flow-variation graphs and effectively improves their genome fraction and NGA50. While the greedy strategies in building candidate paths make VG-Flow tractable, VG-Flow may be forced to use more (incorrect) paths to cover all strains (*i*.*e*. high duplication ratio and error rate) when not all the ground-truth paths have been included in its selected candidate set. VStrains uses contigs, paired-end reads, and coverage information to extract strain-specific paths iteratively from assembly graphs built by SPAdes (VStrains+SPAdes) and achieves the best overall performance on a comprehensive set of metrics (including genome fraction, duplication ratio, NGA50, error rate and number of contigs). It is worth noting that VStrains is able to adapt the general-purpose assembler SPAdes to assemble highly similar strains and even outperform existing specialized strain-aware assemblers.

**Table 2:** Performance of de novo and reference-based approaches on reconstructing viral strains in simulated datasets

| **6 Poliovirus Strains** | Genome Fraction (GF) | Duplication Ratio | NGA50(# ref strains >50% GF) | Error Rate (mis+indel+N's) | # Contigs (> 500 bp) |
| --- | --- | --- | --- | --- | --- |
| PredictHaplo | 16.68% | **1.00** | 7459(1) | 0.871% | 1 |
| PEHaplo | 82.68% | **1.00** | 7393(5) | 0.160% | 5 |
| Haploflow | 61.40% | **1.00** | 6671(3) | 0.517% | 5 |
| viaDBG | 68.80% | 2.51 | 2535(6) | 0.018% | 48 |
| SAVAGE | 85.03% | 1.70 | 2930(5) | **0.014%** | 50 |
| VG-Flow+SAVAGE | 61.69% | 1.27 | 5656(4) | 0.020% | 8 |
| SPAdes | 43.82% | 1.01 | 5706(2) | 0.228% | 8 |
| VG-Flow+SPAdes | - | - | - | - | - |
| VStrains+SPAdes | **89.67%** | **1.00** | **6682(6)** | 0.087% | **6** |
| **10 HCV Strains** | Genome Fraction (GF) | Duplication Ratio | NGA50(# ref strains >50% GF) | Error Rate (mis+indel+N's) | # Contigs (> 500 bp) |
| PredictHaplo | 89.97% | **1.00** | 9292(9) | 0.325% | 9 |
| PEHaplo | 95.95% | 1.01 | 8859(10) | 0.013% | 12 |
| Haploflow | 62.09% | 1.55 | 8893(6) | 3.834% | 22 |
| viaDBG | 97.66% | 2.18 | 9033(10) | **0.002%** | 24 |
| SAVAGE | 99.52% | 1.07 | 9059(10) | **0.002%** | 18 |
| VG-Flow+SAVAGE | **99.67%** | **1.00** | **9264(10)** | 0.003% | **10** |
| SPAdes | 90.81% | **1.00** | 8840(10) | 0.006% | **10** |
| VG-Flow+SPAdes | 94.11% | 1.10 | 8687(10) | 0.016% | 12 |
| VStrains+SPAdes | 98.16% | **1.00** | 9124(10) | 0.046% | **10** |
| **15 ZIKV Strains** | Genome Fraction (GF) | Duplication Ratio | NGA50(# ref strains >50% GF) | Error Rate (mis+indel+N's) | # Contigs (> 500 bp) |
| PredictHaplo | 46.66% | **1.00** | 10269(7) | 0.427% | 7 |
| PEHaplo | 84.24% | 1.5 | 6256(15) | 0.402% | 46 |
| Haploflow | 21.82% | 4.25 | 10198(3) | 4.167% | 26 |
| viaDBG | 93.04% | 2.97 | 5136(15) | 0.028% | 276 |
| SAVAGE | 98.85% | 1.47 | 4031(15) | **0.011%** | 116 |
| VG-Flow+SAVAGE | 98.23% | 1.20 | 10081(15) | 0.077% | 18 |
| SPAdes | 66.71% | **1.00** | 6447(11) | 0.037% | 27 |
| VG-Flow+SPAdes | - | - | - | - | - |
| VStrains+SPAdes | **98.87%** | **1.00** | **10130(15)** | 0.068% | **16** |

### 4.2 Performance on Real Datasets

Table 3 summarizes the performance of VStrains and other de novo and reference-based approaches on two real datasets, 5-HIV-labmix [14] and 2-SARS-COV-2 [37]. While viaDBG results in high genome fraction and low error rate, it comes at a cost of (extremely) high duplication ratio and an excessive number of contigs, making it infeasible to distinguish correct and incorrect contigs from its output. As a reference-based approach, PredictHaplo achieves a high genome fraction (99.22%) and high NGA50 (9604) for 5-HIV-labmix dataset because a ground-truth viral strain was provided as its input. However, in reality, accurate reference genomes are usually not available and species are unknown in viral quasispecies, it is impossible to provide good reference genomes for PredictHaplo.

**Table 3:** Performance of de novo and reference-based approaches on reconstructing viral strains in real datasets

| **5 HIV-labmix Strains** | Genome Fraction (GF) | Duplication Ratio | NGA50(# ref strains >50% GF) | Error Rate (mis+indel+N's) | # Contigs (> 500 bp) |
| --- | --- | --- | --- | --- | --- |
| PredictHaplo | **99.22%** | **1.00** | **9604(5)** | 1.259% | **5** |
| PEHaplo | 84.19% | 1.55 | 1915(5) | 0.300% | 41 |
| Haploflow | 56.80% | 1.26 | 4832(4) | 2.675% | 18 |
| viaDBG | 92.80% | 13.54 | 5942(5) | **0.071%** | 228 |
| SAVAGE | 87.34% | 1.56 | 1397(5) | 0.115% | 72 |
| VG-Flow+SAVAGE | 77.70% | 3.12 | 7235(4) | 1.038% | 23 |
| SPAdes | 49.11% | 1.07 | 614(3) | 0.515% | 33 |
| VG-Flow+SPAdes | 79.75% | 2.27 | 1938(5) | 1.137% | 78 |
| VStrains+SPAdes | 86.42% | 1.30 | 7583(5) | 0.237% | 12 |
| **2 SARS-COV-2 Strains** | Genome Fraction (GF) | Duplication Ratio | NGA50(# ref strains >50% GF) | Error Rate (mis+indel+N's) | # Contigs (> 500 bp) |
| PredictHaplo | 50.02% | 9.00 | 30347(1) | 0.024% | 9 |
| PEHaplo | 67.10% | 1.24 | 21822(1) | 0.073% | 4 |
| Haploflow | 54.78% | 1.12 | 30308(1) | 0.059% | **3** |
| viaDBG | **80.96%** | 1.52 | 5501(2) | **0.004%** | 20 |
| SAVAGE | - | - | - | - | - |
| VG-Flow+SAVAGE | - | - | - | - | - |
| SPAdes | 48.44% | **1.00** | 795(1) | 0.014% | 7 |
| VG-Flow+SPAdes | - | - | - | - | - |
| VStrains+SPAdes | 63.37% | **1.00** | **12272(2)** | 0.013% | **3** |

Similar to the performance on the simulated datasets, SPAdes results in very fragmented assemblies with low genome fraction (49.11%) and NGA50 (614) for 5-HIV-labmix dataset. SAVAGE as a specialized strain-aware assembler produces almost double the genome fraction (87.34%) and NGA50 (1397) compare to SPAdes. While VG-Flow+SAVAGE increases the NGA50 to 7235, it decreases the genome fraction to 77.7%, doubles the duplication ratio (from 1.56 to 3.12), and significantly increase the error rate from 0.115% to 1.038%, which indicates its limitation on handling real datasets. On the other hand, VStrains+SPAdes significantly improves the overall performance on SPAdes, *e*.*g*., increase genome fraction from 49.11% to 86.42%, NGA50 from 614 to 7583, and decreases the error rate from 0.515% to 0.237%. Note that VG-Flow+SPAdes fails to achieve such an improvement as VG-Flow is mainly designed to couple with SAVAGE. However, SAVAGE typically assumes at least 10,000x total coverage of viral sequencing data (quote from its GitHub site), which limits the application of VG-Flow on datasets without ultra-high coverage (e.g., on the 2-SARS-COV-2 dataset with only 4,000x coverage).

## 5 Software and Resource Usage

All the experiments were run under the National Computational Infrastructure (NCI) Gadi supercomputer by submitting jobs to the Gadi biodev queue with the default job dependencies. The allocated RAM size was limited to 500GB and CPU time was limited to 800 hours. The peak memory (maximum resident set size) refers to the peak amount of memory throughout the program execution. Table 4 summarize the CPU time and peak memory for different approaches on all viral quasispecies benchmarks, respectively.

**Table 4:**
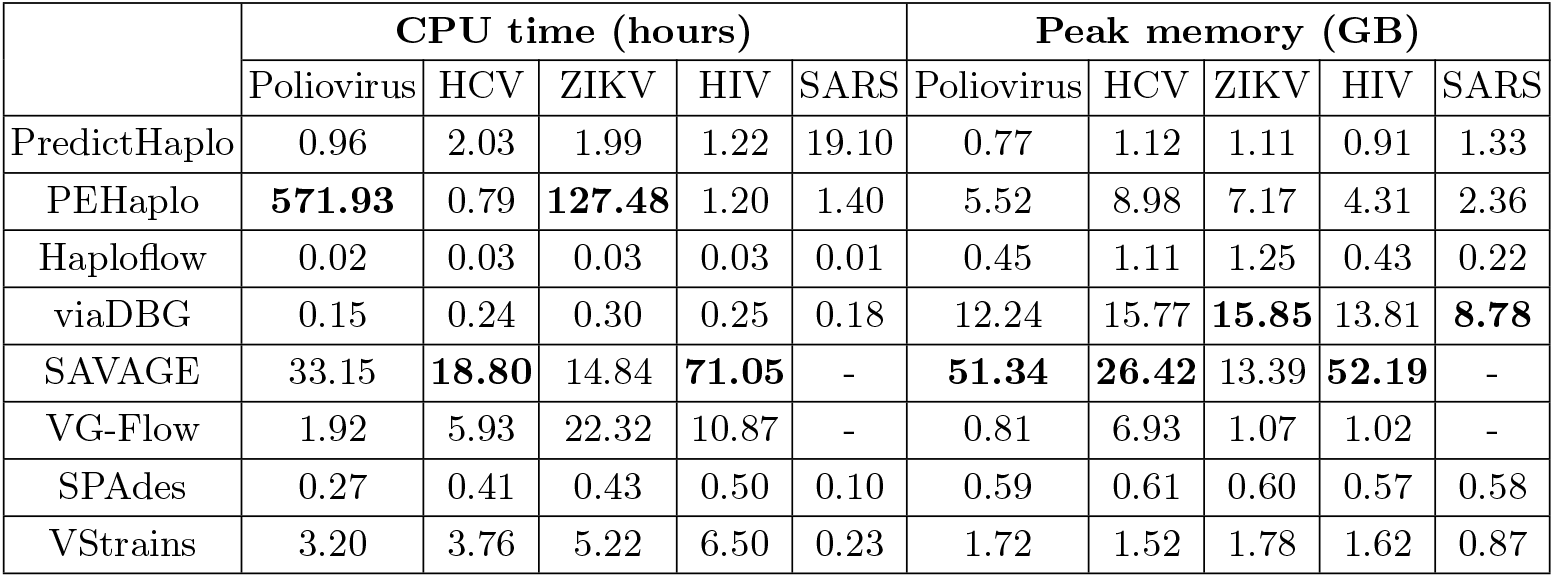
CPU Time and peak memory usage of de novo and reference-based approaches on viral haplotype reconstruction

|  | CPU time (hours) |  |  |  |  | Peak memory (GB) |  |  |  |  |
| --- | --- | --- | --- | --- | --- | --- | --- | --- | --- | --- |
|  | Poliovirus | HCV | ZIKV | HIV | SARS | Poliovirus | HCV | ZIKV | HIV | SARS |
| PredictHaplo | 0.96 | 2.03 | 1.99 | 1.22 | 19.10 | 0.77 | 1.12 | 1.11 | 0.91 | 1.33 |
| PEHaplo | **571.93** | 0.79 | **127.48** | 1.20 | 1.40 | 5.52 | 8.98 | 7.17 | 4.31 | 2.36 |
| Haploflow | 0.02 | 0.03 | 0.03 | 0.03 | 0.01 | 0.45 | 1.11 | 1.25 | 0.43 | 0.22 |
| viaDBG | 0.15 | 0.24 | 0.30 | 0.25 | 0.18 | 12.24 | 15.77 | **15.85** | 13.81 | **8.78** |
| SAVAGE | 33.15 | **18.80** | 14.84 | **71.05** | - | **51.34** | **26.42** | 13.39 | **52.19** | - |
| VG-Flow | 1.92 | 5.93 | 22.32 | 10.87 | - | 0.81 | 6.93 | 1.07 | 1.02 | - |
| SPAdes | 0.27 | 0.41 | 0.43 | 0.50 | 0.10 | 0.59 | 0.61 | 0.60 | 0.57 | 0.58 |
| VStrains | 3.20 | 3.76 | 5.22 | 6.50 | 0.23 | 1.72 | 1.52 | 1.78 | 1.62 | 0.87 |

From Table 4, we observe that Haploflow is much more efficient than all other approaches in runtime and peak memory, but may not be a preferred tool due to its extremely high error rate and low genome fraction (as shown in Table 2 and 3). The general-purpose assembler SPAdes is more efficient than specialized strain-aware assemblers such as PEHaplo, Haploflow, viaDBG and SAVAGE in terms of runtime and peak memory. To reconstruct full-length viral strains from pre-assembled contigs, VG-Flow and VStrains cost comparable running time and memory usage. However, since VG-Flow employs SAVAGE to generate the pre-assembled contigs and the running time and memory usage of SAVAGE are almost the most expensive when compared to other tools, which results in high memory usage and running time of “SAVAGE+VG-Flow". On the contrary, the running time and memory usage of “SPAdes+VStrains” are comparable with other specialized assemblers.

To test the scalability of VStrains, we further compared its runtime and peak memory usage to VG-Flow on simulation datasets with increasing genome size and a variable number of strains (Fig. 4). VG-Flow was selected to compare as it is the most state-of-the-art viral quasispecies assembly post-processing tool. The runtime and memory usage of VStrains do not demonstrate significant correlations with respect to the increase in the number of strains while the runtime of VG-Flow increases with the number of strains. When increasing the genome size, the runtime and memory usage of both VStrain and VG-Flow increased (Fig. 4 (a) and (c)). It is worth noting that, when including the runtime and memory usage of the pre-assemblers (SAVAGE and SPAdes), SAVAGE+VG-Flow is much more sensitive to the genome size than VStrains+SPAdes (Fig. 4 (b) and (d)). Thus, we concluded that VStrains+SPAdes is more efficient than VG-Flow+SAVAGE in the analysis of large-scale datasets. Taking into account the good performance of VStrains+SPAdes at Table 2 and Table 3, it is more cost-effective to employ VStrains as a postprocessing tool for SPAdes in terms of both performance and program efficiency compare to VG-Flow.

**Fig. 4:**
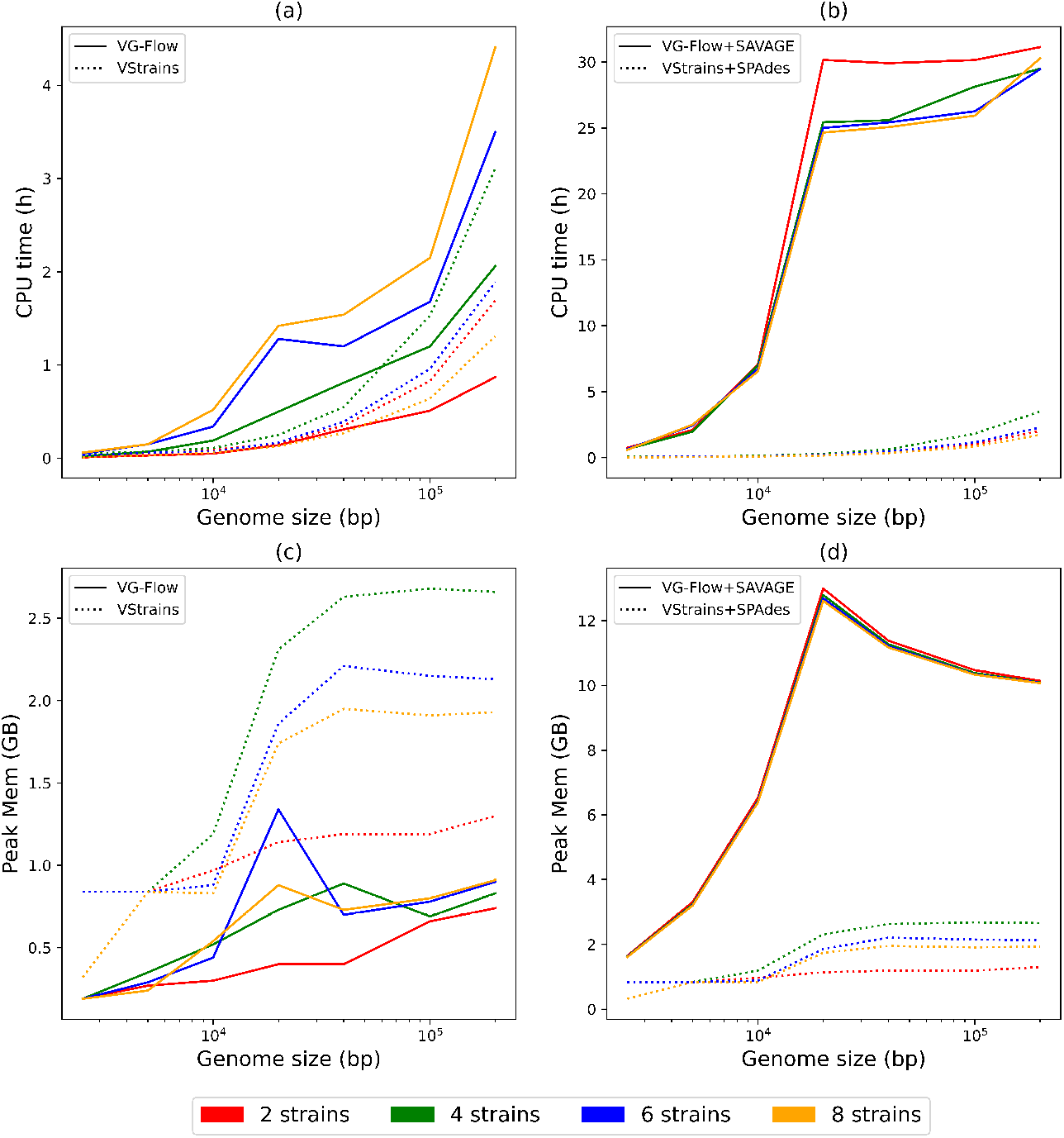
CPU time and peak memory for VG-Flow, VStrains, VG-Flow+SAVAGE, and VStrains+SPAdes on simulated datasets consist of 2, 4, 6, 8 strains with increasing genome size (bp) (2500, 5000, 10.000, 20.000, 40.000, 100.000, 200.000). The x-axis is plotted on a logarithmic scale. The CPU time for VStrains+SPAdes (VG-Flow+SAVAGE) is the addition of VStrains (VG-Flow) and SPAdes (SAVAGE), and the peak memory for VStrains+SPAdes (VG-Flow+SAVAGE) is the maximum between VStrains (VG-Flow) and SPAdes (SAVAGE).

## 6 Conclusion and Discussion

In VStrains, we first introduce a strategy to canonize the strandedness of assembly graphs from SPAdes, which reduces the strain reconstruction problem into the path extraction problem. Secondly, we use all available *k*-mer pairs in paired-end reads to infer *PE* links in the assembly graph produced by SPAdes. Thirdly, we propose an effective way to incorporate *PE* links together with contigs and coverage information to disentangle the assembly graphs. Finally, we demonstrate how to extract confident strain-specific paths via iterative contig-based path extraction. Experimental results on both simulated and real datasets show that VStrains achieves the best overall performance among the state-of-the-art approaches.

Currently, VStrains relies on both assembly graphs and contigs from SPAdes and thus cannot couple with assemblers which does not explicitly output assembly graphs such as SAVAGE. The current implementation of VStrains requires additional alignments of paired-end reads to vertices in assembly graphs to infer *PE* links, which dominates the total runtime and peak memory usage. It is worth exploring how to make VStrains more flexible and efficient.

With the advance of third-generation sequencing (TGS), multiple approaches (including Strainline [22], Strainberry [35], VirStrain [21], viralFlye [2], *etc*.) have been proposed for strain-aware assembly using TGS data. VStrains has the potential to be extended to handle TGS data by taking advantages of assembly graphs built from Canu [19], Flye [18], wtdbg [32] and others.

## Acknowledgements

We want to thank the anonymous reviewers for providing valuable and detailed feedback. We thank Hansheng Xue and Vijini Mallawaarachchi for testing the machine learning-based method (GAEseq, CAECSeq, and NeurHap) and SPAdes-series assembler (MetaSPAdes and MetaviralSPAdes). We thank Lianrong Pu for polishing the paper and providing valuable advice to us. This research was undertaken with the assistance of resources and services from the National Computational Infrastructure (NCI), which is supported by the Australian Government.

