## Supplementary Material for "VStrains: De Novo Reconstruction of Viral Strains via Iterative Path Extraction From Assembly Graphs"

Runpeng Luo[0000−0002−4413−9122] and Yu Lin(✉)[0000−0001−6339−2644]

School of Computing, Australian National University, Canberra, Australia
{john.luo,yu.lin}@anu.edu.au

#### S1. Comparison among SPAdes-series assemblers

In consistent with previous observations [3,8], Table S1.1 demonstrate that SPAdes [4] in general outperforms other more specific SPAdes-series assemblers such as metaSPAdes [17], MetaviralSPAdes [1], rnaviralSPAdes [5] and coronaSPAdes [15], especially in terms of duplication ratio and error rates. Therefore, VStrains employs the general-purpose assembler SPAdes to build assembly graphs.

**Table S1.1.** Comparison among SPAdes-series assemblers on real datasets

| 5 HIV-labmix Strains | Genome Fraction (GF) | Duplication Ratio | NGA50(# ref strains >50% GF) | Error Rate (mis+indel+N's) | # contigs > 500 bp |
|---|---|---|---|---|---|
| SPAdes | 49.11% | 1.07 | 614(3) | 0.515% | 33 |
| MetaSPAdes | 40.61% | 3.25 | 3990(3) | 1.859% | 20 |
| MetaviralSPAdes | - | - | - | - | - |
| rnaviralSPAdes | 39.65% | 1.06 | 797.5(2) | 0.367% | 15 |

| 2 SARS-COV-2 Strains | Genome Fraction (GF) | Duplication Ratio | NGA50(# ref strains >50% GF) | Error Rate (mis+indel+N's) | # contigs > 500 bp |
|---|---|---|---|---|---|
| SPAdes | 48.44% | 1.00 | 795(1) | 0.014% | 7 |
| MetaSPAdes | 49.98% | 3.30 | 30314(1) | 0.036% | 1 |
| MetaviralSPAdes | 49.97% | 3.30 | 30309(1) | 0.036% | 1 |
| rnaviralSPAdes | 49.98% | 3.30 | 30314(1) | 0.033% | 1 |
| coronaSPAdes | 49.98% | 3.30 | 30314(1) | 0.033% | 1 |

Comparing SPAdes, MetaSPAdes, MetaviralSPAdes, rnaviralSPAdes and coronaSPAdes on two real datasets, 5 HIV-labmix and 2 SARS-COV-2. MetaviralSPAdes failed to output any contigs for 5 HIV-labmix dataset.

### S2. Detailed strain-level MetaQUAST evaluation results

The following tables consist of detailed evaluation result from MetaQUAST [16] (–unique-mapping option), specified to individual viral strain for all datasets.

For each assembly, we report the genome fraction, duplication ratio, NGA50, error rate, and number of contigs. Genome fraction is defined as the total number of aligned bases in the reference, divided by the genome size. A base in the reference genome is counted as aligned if there is at least one contig with at least one alignment to this base. Duplication ratio is defined as the total number of aligned bases in the assembly, divided by the total number of aligned bases in the reference. NG50 is the contig length such that using longer or equal length contigs produce half of the bases of the reference genome, whereas NGA50 counts the lengths of aligned blocks instead of contig lengths, such that the contig is broken into smaller pieces when it has a misassembly with respect to the reference genome. Error rate is defined as the sum of mismatch rate, indel rate, and N's rate, which reflects the number of errors with respect to the reference genome size.

Table S2.1.1 to Table S2.1.4 describe the detailed strain-level performance for 6-Poliovirus simulated dataset, Table S2.2.1 to Table S2.2.4 describe the detailed strain-level performance for 10-HCV simulated dataset, Table S2.3.1 to Table S2.3.4 describe the detailed strain-level performance for 15-ZIKV simulated dataset, Table S2.4.1 to Table S2.4.4 describe the detailed strain-level performance for 5-HIV-labmix real dataset, and Table S2.5.1 to Table S2.5.4 describe the detailed strain-level performance for 2-SARS-COV-2 real dataset. Note that S$i$ shown in the Table S2.1.1 to Table S2.3.4 represents the $i$th reference viral strain for the corresponding dataset.

**Table S2.1.1.** Genome fraction on 6-Poliovirus simulated dataset

| | | 6 Poliovirus | | | | | |
|---|---|---|---|---|---|---|---|
| | | S1 | S2 | S3 | S4 | S5 | S6 |
| SPAdes | Genome Fraction(GF)(%) | 24.09 | 25.01 | 100.00 | 88.44 | 11.73 | 13.52 |
| SAVAGE | | 99.69 | 92.45 | 99.50 | 96.64 | 88.32 | 33.61 |
| PEHaplo | | 100.00 | 96.51 | 100.00 | 0.00 | 99.77 | 99.91 |
| viaDBG | | 53.02 | 60.53 | 99.69 | 48.25 | 63.89 | 87.35 |
| Haploflow | | 100.00 | 98.67 | 71.97 | - | 87.62 | 10.30 |
| PredictHaplo | | - | - | - | 100.00 | - | - |
| VG-Flow+SAVAGE | | 73.74 | 99.79 | 99.49 | 52.05 | 32.99 | 12.11 |
| VG-Flow+SPAdes | | - | - | - | - | - | - |
| VStrains+SPAdes | | 100.00 | 90.84 | 100.00 | 97.24 | 90.51 | 59.46 |

**Table S2.1.2.** Duplication ratio on 6-Poliovirus simulated dataset

| | | 6 Poliovirus | | | | | |
|---|---|---|---|---|---|---|---|
| | | S1 | S2 | S3 | S4 | S5 | S6 |
| SPAdes | Duplication Ratio | 1.07 | 1.00 | 1.00 | 1.00 | 1.00 | 1.00 |
| SAVAGE | | 1.95 | 1.89 | 1.00 | 1.88 | 2.01 | 1.17 |
| PEHaplo | | 1.00 | 1.00 | 1.00 | - | 1.00 | 1.00 |
| viaDBG | | 1.65 | 2.99 | 3.95 | 1.42 | 1.65 | 2.28 |
| Haploflow | | 1.00 | 1.00 | 1.00 | - | 1.00 | 1.00 |
| PredictHaplo | | - | - | - | 1.00 | - | - |
| VG-Flow+SAVAGE | | 1.00 | 1.99 | 1.00 | 1.00 | 1.00 | 1.00 |
| VG-Flow+SPAdes | | - | - | - | - | - | - |
| VStrains+SPAdes | | 1.00 | 1.00 | 1.00 | 1.00 | 1.00 | 1.00 |

**Table S2.1.3.** NGA50 on 6-Poliovirus simulated dataset

| | | 6 Poliovirus | | | | | |
|---|---|---|---|---|---|---|---|
| | | S1 | S2 | S3 | S4 | S5 | S6 |
| SPAdes | NGA50 | - | - | 7460 | 3951 | - | - |
| SAVAGE | | 1453 | 2194 | 7423 | 2188 | 1394 | - |
| PEHaplo | | 7428 | 7192 | 7460 | - | 7440 | 7447 |
| viaDBG | | 815 | 3067 | 7437 | 962 | 869 | 2060 |
| Haploflow | | 7428 | 7353 | 5369 | - | 6534 | - |
| PredictHaplo | | - | - | - | 7459 | - | - |
| VG-Flow+SAVAGE | | 3882 | 7436 | 7422 | 3882 | - | - |
| VG-Flow+SPAdes | | - | - | - | - | - | - |
| VStrains+SPAdes | | 7428 | 6769 | 7460 | 7253 | 6749 | 4432 |

**Table S2.1.4.** Error rate (N's + mismatches + indels) on 6-Poliovirus simulated dataset

| | | 6 Poliovirus | | | | | |
|---|---|---|---|---|---|---|---|
| | | S1 | S2 | S3 | S4 | S5 | S6 |
| SPAdes | Error Rate (%) | 0.000 | 0.000 | 0.268 | 0.197 | 1.371 | 0.000 |
| SAVAGE | | 0.007 | 0.031 | 0.000 | 0.007 | 0.023 | 0.000 |
| PEHaplo | | 0.148 | 0.083 | 0.000 | - | 0.323 | 0.242 |
| viaDBG | | 0.015 | 0.022 | 0.000 | 0.039 | 0.000 | 0.054 |
| Haploflow | | 0.000 | 0.598 | 0.671 | - | 0.949 | 0.000 |
| PredictHaplo | | - | - | - | 0.871 | - | - |
| VG-Flow+SAVAGE | | 0.018 | 0.040 | 0.000 | 0.000 | 0.000 | 0.000 |
| VG-Flow+SPAdes | | - | - | - | - | - | - |
| VStrains+SPAdes | | 0.000 | 0.000 | 0.040 | 0.193 | 0.119 | 0.226 |

**Table S2.2.1.** Genome fraction on 10-HCV simulated dataset

| | | 10 HCV | | | | | | | | | |
|---|---|---|---|---|---|---|---|---|---|---|---|
| | | S1 | S2 | S3 | S4 | S5 | S6 | S7 | S8 | S9 | S10 |
| SPAdes | Genome Fraction(GF)(%) | 97.02 | 80.84 | 87.42 | 100.00 | 96.37 | 84.39 | 84.34 | 86.76 | 95.76 | 95.18 |
| SAVAGE | | 99.73 | 99.32 | 99.56 | 99.52 | 99.60 | 98.97 | 99.73 | 99.58 | 99.40 | 99.79 |
| PEHaplo | | 100.00 | 92.64 | 89.63 | 100.00 | 90.03 | 96.67 | 100.00 | 100.00 | 90.94 | 99.56 |
| viaDBG | | 99.90 | 97.16 | 94.95 | 97.74 | 98.72 | 94.57 | 99.29 | 98.59 | 95.93 | 99.70 |
| Haploflow | | 6.04 | 15.96 | 94.61 | 95.73 | 93.90 | 91.00 | 98.96 | 9.25 | 15.24 | 99.98 |
| PredictHaplo | | 100.00 | 100.00 | - | 99.94 | 99.97 | 100.00 | 99.84 | 99.94 | 99.99 | 99.84 |
| VG-Flow+SAVAGE | | 99.82 | 99.56 | 99.64 | 99.82 | 99.72 | 99.32 | 99.72 | 99.82 | 99.56 | 99.72 |
| VG-Flow+SPAdes | | 100.00 | 84.26 | 90.90 | 100.00 | 99.99 | 84.73 | 92.48 | 93.46 | 100.00 | 95.25 |
| VStrains+SPAdes | | 98.93 | 98.33 | 98.33 | 100.00 | 98.66 | 97.77 | 98.28 | 95.14 | 97.67 | 98.53 |

**Table S2.2.2.** Duplication ratio on 10-HCV simulated dataset

| | | 10 HCV | | | | | | | | | |
|---|---|---|---|---|---|---|---|---|---|---|---|
| | | S1 | S2 | S3 | S4 | S5 | S6 | S7 | S8 | S9 | S10 |
| SPAdes | Duplication Ratio | 1.00 | 1.00 | 1.00 | 1.00 | 1.00 | 1.00 | 1.00 | 1.00 | 1.00 | 1.00 |
| SAVAGE | | 1.00 | 1.07 | 1.11 | 1.00 | 1.07 | 1.30 | 1.00 | 1.00 | 1.10 | 1.06 |
| PEHaplo | | 1.00 | 1.00 | 1.00 | 1.00 | 1.00 | 1.00 | 1.06 | 1.00 | 1.00 | 1.00 |
| viaDBG | | 2.96 | 1.99 | 1.96 | 1.00 | 2.00 | 1.94 | 1.00 | 2.97 | 2.98 | 2.99 |
| Haploflow | | 1.00 | 1.00 | 2.11 | 2.05 | 1.27 | 1.69 | 1.00 | 1.00 | 1.00 | 1.47 |
| PredictHaplo | | 1.00 | 1.00 | - | 1.00 | 1.00 | 1.00 | 1.00 | 1.00 | 1.00 | 1.00 |
| VG-Flow+SAVAGE | | 1.00 | 1.00 | 1.00 | 1.00 | 1.00 | 1.00 | 1.00 | 1.00 | 1.00 | 1.00 |
| VG-Flow+SPAdes | | 1.00 | 1.00 | 1.98 | 1.00 | 1.00 | 1.00 | 1.01 | 1.00 | 1.00 | 1.00 |
| VStrains+SPAdes | | 1.00 | 1.00 | 1.00 | 1.00 | 1.00 | 1.00 | 1.00 | 1.00 | 1.00 | 1.00 |

**Table S2.2.3.** NGA50 on 10-HCV simulated dataset

| | | 10 HCV | | | | | | | | | |
|---|---|---|---|---|---|---|---|---|---|---|---|
| | | S1 | S2 | S3 | S4 | S5 | S6 | S7 | S8 | S9 | S10 |
| SPAdes | NGA50 | 8997 | 7505 | 8106 | 9302 | 8960 | 7845 | 7853 | 8070 | 8903 | 8862 |
| SAVAGE | | 9248 | 9221 | 9148 | 9257 | 8939 | 8174 | 9286 | 9263 | 8800 | 9251 |
| PEHaplo | | 9273 | 8601 | 8311 | 9302 | 8371 | 8986 | 9311 | 9302 | 8455 | 8674 |
| viaDBG | | 9264 | 9020 | 8805 | 9092 | 9143 | 8593 | 9245 | 8967 | 8918 | 9283 |
| Haploflow | | - | - | 8750 | 8897 | 8731 | 8459 | 9214 | - | - | 9309 |
| PredictHaplo | | 9273 | 9284 | - | 9296 | 9295 | 9296 | 9296 | 9296 | 9296 | 9296 |
| VG-Flow+SAVAGE | | 9256 | 9243 | 9240 | 9285 | 9272 | 9233 | 9285 | 9285 | 9256 | 9285 |
| VG-Flow+SPAdes | | 9273 | 7823 | 8429 | 9302 | 9297 | 7876 | 8010 | 8694 | 9297 | 8869 |
| VStrains+SPAdes | | 9174 | 9129 | 9118 | 9302 | 9173 | 9089 | 9151 | 8850 | 9080 | 9174 |

**Table S2.2.4.** Error rate (N's + mismatches + indels) on 10-HCV simulated dataset

| | | 10 HCV | | | | | | | | | |
|---|---|---|---|---|---|---|---|---|---|---|---|
| | | S1 | S2 | S3 | S4 | S5 | S6 | S7 | S8 | S9 | S10 |
| SPAdes | Error Rate (%) | 0.000 | 0.000 | 0.000 | 0.000 | 0.045 | 0.000 | 0.000 | 0.000 | 0.011 | 0.000 |
| SAVAGE | | 0.000 | 0.020 | 0.000 | 0.000 | 0.000 | 0.000 | 0.000 | 0.000 | 0.000 | 0.000 |
| PEHaplo | | 0.000 | 0.035 | 0.000 | 0.000 | 0.012 | 0.000 | 0.040 | 0.011 | 0.000 | 0.043 |
| viaDBG | | 0.000 | 0.000 | 0.017 | 0.000 | 0.000 | 0.000 | 0.000 | 0.000 | 0.000 | 0.004 |
| Haploflow | | 3.571 | 2.699 | 3.969 | 3.742 | 3.938 | 4.524 | 3.071 | 2.442 | 2.682 | 3.821 |
| PredictHaplo | | 0.000 | 0.086 | - | 0.011 | 0.000 | 0.000 | 0.839 | 0.237 | 0.581 | 1.173 |
| VG-Flow+SAVAGE | | 0.000 | 0.022 | 0.000 | 0.000 | 0.000 | 0.000 | 0.000 | 0.011 | 0.000 | 0.000 |
| VG-Flow+SPAdes | | 0.011 | 0.013 | 0.006 | 0.000 | 0.065 | 0.000 | 0.012 | 0.023 | 0.032 | 0.000 |
| VStrains+SPAdes | | 0.000 | 0.153 | 0.033 | 0.000 | 0.000 | 0.154 | 0.098 | 0.011 | 0.000 | 0.011 |

**Table S2.3.1.** Genome fraction on 15-ZIKV simulated dataset

| | | 15 ZIKV | | | | | | | | | | | | | | |
|---|---|---|---|---|---|---|---|---|---|---|---|---|---|---|---|---|
| | | S1 | S2 | S3 | S4 | S5 | S6 | S7 | S8 | S9 | S10 | S11 | S12 | S13 | S14 | S15 |
| SPAdes | Genome Fraction(GF)(%) | - | 17.26 | 11.02 | 62.76 | 27.13 | 63.72 | 57.73 | 86.20 | 83.80 | 92.33 | 100.00 | 99.99 | 98.60 | 100.00 | 100.00 |
| SAVAGE | | 94.90 | 98.18 | 97.14 | 98.92 | 98.81 | 99.38 | 99.35 | 99.24 | 99.35 | 99.39 | 99.58 | 99.45 | 99.68 | 99.73 | 99.71 |
| PEHaplo | | 76.22 | 99.98 | 84.54 | 99.98 | 75.12 | 51.81 | 53.34 | 75.63 | 94.83 | 100.00 | 99.97 | 99.99 | 77.95 | 74.21 | 100.00 |
| viaDBG | | 74.74 | 83.57 | 69.18 | 95.30 | 96.23 | 99.67 | 95.95 | 97.73 | 97.53 | 98.14 | 99.32 | 99.91 | 89.05 | 99.35 | 99.96 |
| Haploflow | | 99.97 | 99.07 | 99.26 | - | - | - | - | - | - | - | - | - | 9.86 | 6.63 | 12.44 |
| PredictHaplo | | - | 99.97 | - | 100.00 | - | - | - | - | - | 100.00 | - | 100.00 | 100.00 | 100.00 | 100.00 |
| VG-Flow+SAVAGE | | 92.28 | 98.27 | 99.32 | 92.31 | 98.80 | 98.58 | 98.72 | 99.15 | 98.58 | 99.53 | 99.54 | 99.50 | 99.60 | 99.62 | 99.60 |
| VG-Flow+SPAdes | | - | - | - | - | - | - | - | - | - | - | - | - | - | - | - |
| VStrains+SPAdes | | 97.46 | 99.09 | 97.20 | 99.49 | 99.09 | 97.20 | 95.12 | 98.58 | 100.00 | 99.89 | 100.00 | 99.99 | 100.00 | 100.00 | 100.00 |

**Table S2.3.2.** Duplication ratio on 15-ZIKV simulated dataset

| | | 15 ZIKV | | | | | | | | | | | | | | |
|---|---|---|---|---|---|---|---|---|---|---|---|---|---|---|---|---|
| | | S1 | S2 | S3 | S4 | S5 | S6 | S7 | S8 | S9 | S10 | S11 | S12 | S13 | S14 | S15 |
| SPAdes | Duplication Ratio | - | 1.00 | 1.00 | 1.00 | 1.00 | 1.02 | 1.02 | 1.00 | 1.00 | 1.01 | 1.00 | 1.00 | 1.00 | 1.00 | 1.00 |
| SAVAGE | | 1.37 | 1.51 | 1.45 | 2.05 | 1.50 | 1.74 | 1.71 | 1.76 | 2.08 | 1.18 | 1.00 | 1.11 | 1.16 | 1.18 | 1.24 |
| PEHaplo | | 1.85 | 2.05 | 1.00 | 1.75 | 1.08 | 1.00 | 1.76 | 1.25 | 1.22 | 2.90 | 1.00 | 1.00 | 1.95 | 1.00 | 1.37 |
| viaDBG | | 1.50 | 2.08 | 1.90 | 2.35 | 3.61 | 3.39 | 3.20 | 2.43 | 3.52 | 3.65 | 4.01 | 3.58 | 1.56 | 4.13 | 2.63 |
| Haploflow | | 6.06 | 4.44 | 3.20 | - | - | - | - | - | - | - | - | - | 1.00 | 1.00 | 1.00 |
| PredictHaplo | | - | 1.00 | - | 1.00 | - | - | - | - | - | 1.00 | - | 1.00 | 1.00 | 1.00 | 1.00 |
| VG-Flow+SAVAGE | | 1.00 | 1.00 | 1.00 | 2.00 | 1.00 | 1.00 | 1.00 | 2.00 | 1.99 | 1.00 | 1.00 | 1.00 | 1.00 | 1.00 | 1.00 |
| VG-Flow+SPAdes | | - | - | - | - | - | - | - | - | - | - | - | - | - | - | - |
| VStrains+SPAdes | | 1.00 | 1.00 | 1.00 | 1.00 | 1.03 | 1.00 | 1.00 | 1.00 | 1.00 | 1.00 | 1.00 | 1.00 | 1.00 | 1.00 | 1.00 |

**Table S2.3.3.** NGA50 on 15-ZIKV simulated dataset

| | | 15 ZIKV | | | | | | | | | | | | | | |
|---|---|---|---|---|---|---|---|---|---|---|---|---|---|---|---|---|
| | | S1 | S2 | S3 | S4 | S5 | S6 | S7 | S8 | S9 | S10 | S11 | S12 | S13 | S14 | S15 |
| SPAdes | NGA50 | - | - | - | 696 | - | 1479 | 844 | 8185 | 3316 | 5219 | 10269 | 10268 | 10107 | 10269 | 10269 |
| SAVAGE | | 1263 | 1541 | 1774 | 2201 | 3253 | 2192 | 3588 | 3077 | 2580 | 4083 | 10226 | 6345 | 6269 | 6322 | 5749 |
| PEHaplo | | 7813 | 8157 | 3255 | 6383 | 3240 | 1353 | 3812 | 2540 | 6488 | 10251 | 10266 | 10268 | 5808 | 7621 | 6590 |
| viaDBG | | 888 | 825 | 1072 | 2188 | 4252 | 2370 | 6451 | 1954 | 3466 | 10060 | 10197 | 10260 | 6910 | 10201 | 5940 |
| Haploflow | | 10266 | 10172 | 10156 | - | - | - | - | - | - | - | - | - | - | - | - |
| PredictHaplo | | - | 10266 | - | 10269 | - | - | - | - | - | 10269 | - | 10269 | 10269 | 10269 | 10269 |
| VG-Flow+SAVAGE | | 9460 | 10090 | 10199 | 9463 | 10145 | 10123 | 10120 | 10181 | 10123 | 10203 | 10222 | 10218 | 10210 | 10230 | 10228 |
| VG-Flow+SPAdes | | - | - | - | - | - | - | - | - | - | - | - | - | - | - | - |
| VStrains+SPAdes | | 9991 | 10175 | 9981 | 10199 | 9907 | 9981 | 9751 | 10123 | 10269 | 10240 | 10269 | 10268 | 10251 | 10269 | 10269 |

**Table S2.3.4.** Error rate (N's + mismatches + indels) on 15-ZIKV simulated dataset

| | | 15 ZIKV | | | | | | | | | | | | | | |
|---|---|---|---|---|---|---|---|---|---|---|---|---|---|---|---|---|
| | | S1 | S2 | S3 | S4 | S5 | S6 | S7 | S8 | S9 | S10 | S11 | S12 | S13 | S14 | S15 |
| SPAdes | Error Rate (%) | - | 0.451 | 0.265 | 0.249 | 0.000 | 0.000 | 0.033 | 0.011 | 0.035 | 0.000 | 0.010 | 0.000 | 0.030 | 0.010 | 0.000 |
| SAVAGE | | 0.022 | 0.013 | 0.048 | 0.019 | 0.000 | 0.006 | 0.012 | 0.006 | 0.009 | 0.000 | 0.010 | 0.000 | 0.000 | 0.008 | 0.000 |
| PEHaplo | | 0.831 | 0.737 | 0.622 | 0.496 | 0.252 | 0.094 | 0.083 | 0.113 | 0.437 | 0.593 | 0.000 | 0.000 | 0.244 | 0.092 | 0.342 |
| viaDBG | | 0.017 | 0.045 | 0.037 | 0.009 | 0.084 | 0.017 | 0.076 | 0.033 | 0.014 | 0.005 | 0.000 | 0.000 | 0.141 | 0.000 | 0.030 |
| Haploflow | | 4.113 | 4.239 | 4.323 | - | - | - | - | - | - | - | - | - | 3.066 | 1.322 | 2.584 |
| PredictHaplo | | - | 0.955 | - | 0.516 | - | - | - | - | - | 0.234 | - | 0.886 | 0.010 | 0.010 | 0.380 |
| VG-Flow+SAVAGE | | 0.264 | 0.020 | 0.294 | 0.122 | 0.000 | 0.148 | 0.059 | 0.020 | 0.159 | 0.000 | 0.010 | 0.000 | 0.000 | 0.010 | 0.000 |
| VG-Flow+SPAdes | | - | - | - | - | - | - | - | - | - | - | - | - | - | - | - |
| VStrains+SPAdes | | 0.130 | 0.079 | 0.030 | 0.265 | 0.172 | 0.120 | 0.041 | 0.040 | 0.010 | 0.049 | 0.000 | 0.000 | 0.078 | 0.010 | 0.000 |

**Table S2.4.1.** Genome fraction on 5-HIV-labmix real dataset

| | | 5 HIV-labmix | | | | |
|---|---|---|---|---|---|---|
| | | 89.6 | HXB2 | JRCSF | NL43 | YU2 |
| SPAdes | Genome Fraction(GF)(%) | 59.47 | 36.09 | 44.04 | 49.43 | 56.42 |
| SAVAGE | | 94.09 | 71.64 | 95.44 | 94.05 | 81.64 |
| PEHaplo | | 85.86 | 78.12 | 91.58 | 85.98 | 79.53 |
| viaDBG | | 94.01 | 88.71 | 94.79 | 93.81 | 92.70 |
| Haploflow | | 53.15 | 12.21 | 95.44 | 30.42 | 93.51 |
| PredictHaplo | | 98.92 | 99.18 | 99.79 | 99.10 | 99.10 |
| VG-Flow+SAVAGE | | 94.09 | 64.26 | 95.44 | 94.10 | 40.92 |
| VG-Flow+SPAdes | | 88.63 | 40.60 | 87.78 | 93.24 | 88.67 |
| VStrains+SPAdes | | 93.46 | 92.76 | 93.47 | 78.23 | 74.32 |

**Table S2.4.2.** Duplication ratio on 5-HIV-labmix real dataset

| | | 5 HIV-labmix | | | | |
|---|---|---|---|---|---|---|
| | | 89.6 | HXB2 | JRCSF | NL43 | YU2 |
| SPAdes | Duplication Ratio | 1.09 | 1.02 | 1.02 | 1.16 | 1.05 |
| SAVAGE | | 1.35 | 1.23 | 1.76 | 2.11 | 1.24 |
| PEHaplo | | 1.47 | 1.10 | 1.78 | 2.22 | 1.07 |
| viaDBG | | 22.46 | 6.75 | 12.98 | 19.46 | 5.54 |
| Haploflow | | 1.11 | 1.00 | 1.23 | 1.72 | 1.27 |
| PredictHaplo | | 1.00 | 1.00 | 1.00 | 1.00 | 1.00 |
| VG-Flow+SAVAGE | | 2.20 | 1.24 | 4.47 | 4.90 | 1.00 |
| VG-Flow+SPAdes | | 2.47 | 1.84 | 2.16 | 2.43 | 2.21 |
| VStrains+SPAdes | | 1.53 | 1.25 | 1.44 | 1.21 | 1.00 |

**Table S2.4.3.** NGA50 on 5-HIV-labmix real dataset

| | | 5 HIV-labmix | | | | |
|---|---|---|---|---|---|---|
| | | 89.6 | HXB2 | JRCSF | NL43 | YU2 |
| SPAdes | NGA50 | 565 | - | - | 613 | 664 |
| SAVAGE | | 2308 | 751 | 1199 | 1065 | 1662 |
| PEHaplo | | 2163 | 1029 | 2897 | 2685 | 803 |
| viaDBG | | 9129 | 2551 | 6637 | 9108 | 2284 |
| Haploflow | | 550 | - | 9119 | 524 | 9136 |
| PredictHaplo | | 9616 | 9639 | 9500 | 9632 | 9632 |
| VG-Flow+SAVAGE | | 9138 | 1592 | 9063 | 9146 | - |
| VG-Flow+SPAdes | | 1394 | 1210 | 1122 | 1871 | 4091 |
| VStrains+SPAdes | | 8829 | 9001 | 8913 | 5659 | 5512 |

**Table S2.4.4.** Error rate (N's + mismatches + indels) on 5-HIV-labmix real dataset

| | | 5 HIV-labmix | | | | |
|---|---|---|---|---|---|---|
| | | 89.6 | HXB2 | JRCSF | NL43 | YU2 |
| SPAdes | Error Rate (%) | 0.319 | 0.615 | 0.303 | 1.002 | 0.349 |
| SAVAGE | | 0.129 | 0.012 | 0.137 | 0.171 | 0.041 |
| PEHaplo | | 0.286 | 0.262 | 0.398 | 0.281 | 0.219 |
| viaDBG | | 0.041 | 0.074 | 0.113 | 0.066 | 0.100 |
| Haploflow | | 2.867 | 1.855 | 2.395 | 1.945 | 3.261 |
| PredictHaplo | | 1.050 | 1.867 | 0.878 | 0.374 | 2.118 |
| VG-Flow+SAVAGE | | 0.757 | 0.915 | 1.311 | 1.026 | 0.050 |
| VG-Flow+SPAdes | | 1.372 | 0.605 | 1.031 | 1.228 | 1.076 |
| VStrains+SPAdes | | 0.072 | 0.755 | 0.039 | 0.163 | 0.194 |

**Table S2.5.1.** Genome fraction on 2-SARS-COV-2 real dataset

| | | 2 SARS-COV-2 | |
|---|---|---|---|
| | | BA.1 | B.1.1 |
| SPAdes | Genome Fraction(GF)(%) | 52.01 | 44.87 |
| SAVAGE | | - | - |
| PEHaplo | | 34.28 | 99.99 |
| viaDBG | | 70.65 | 91.27 |
| Haploflow | | 9.62 | 99.98 |
| PredictHaplo | | 100.00 | - |
| VG-Flow+SAVAGE | | - | - |
| VG-Flow+SPAdes | | - | - |
| VStrains+SPAdes | | 72.46 | 54.27 |

**Table S2.5.2.** Duplication ratio on 2-SARS-COV-2 real dataset

| | | 2 SARS-COV-2 | |
|---|---|---|---|
| | | BA.1 | B.1.1 |
| SPAdes | Duplication Ratio | 1.00 | 1.00 |
| SAVAGE | | - | - |
| PEHaplo | | 1.00 | 1.32 |
| viaDBG | | 1.69 | 1.38 |
| Haploflow | | 0.99 | 1.13 |
| PredictHaplo | | 9.00 | - |
| VG-Flow+SAVAGE | | - | - |
| VG-Flow+SPAdes | | - | - |
| VStrains+SPAdes | | 1.00 | 1.00 |

**Table S2.5.3.** NGA50 on 2-SARS-COV-2 real dataset

| | | 2 SARS-COV-2 | |
|---|---|---|---|
| | | BA.1 | B.1.1 |
| SPAdes | NGA50 | 795 | - |
| SAVAGE | | - | - |
| PEHaplo | | - | 21822 |
| viaDBG | | 4197 | 6806 |
| Haploflow | | - | 30308 |
| PredictHaplo | | 30347 | - |
| VG-Flow+SAVAGE | | - | - |
| VG-Flow+SPAdes | | - | - |
| VStrains+SPAdes | | 8129 | 16415 |

**Table S2.5.4.** Error rate (N's + mismatches + indels) on 2-SARS-COV-2 real dataset

| | | 2 SARS-COV-2 | |
|---|---|---|---|
| | | BA.1 | B.1.1 |
| SPAdes | Error Rate (%) | 0.000 | 0.029 |
| SAVAGE | | - | - |
| PEHaplo | | 0.202 | 0.031 |
| viaDBG | | 0.008 | 0.000 |
| Haploflow | | 0.380 | 0.032 |
| PredictHaplo | | 0.024 | - |
| VG-Flow+SAVAGE | | - | - |
| VG-Flow+SPAdes | | - | - |
| VStrains+SPAdes | | 0.000 | 0.030 |

### S3. Command lines

For benchmarking purposes, we made use of several existing tools, listed below. All tools under evaluation use default settings unless specified otherwise.

##### Software versions and sources

- Utilities
  - fastp [7]: 0.23.2, https://github.com/OpenGene/fastp
  - BWA [12]: 0.7.17-r1198-dirty, https://github.com/lh3/bwa
  - SAMtools [14]: 1.14-28-g84dfab2 (using htslib 1.14), https://github.com/samtools/samtools
  - vg [10]: v1.40.0 "Suardi", https://github.com/vgteam/vg
  - Gurobi [11]: 9.5.0, https://www.gurobi.com
  - minimap2 [13]: 2.23-r1111, https://github.com/lh3/minimap2
  - MetaQUAST [16]: v5.2.0, https://github.com/ablab/quast
- Assembly tools
  - SPAdes-series [1,4,5,15,17]: v3.15.4, https://github.com/ablab/spades
  - SAVAGE [2]: v0.4.2, https://github.com/HaploConduct/HaploConduct/tree/master/savage
  - VG-Flow [3]: v0.0.4, https://bitbucket.org/jbaaijens/vg-flow/src/master/
  - PredictHaplo [18]: v0.6 paired-end read, https://bmda.dmi.unibas.ch/software.html
  - viaDBG [8]: 1.0, https://github.com/borjaf696/viaDBG
  - Haploflow [9]: 1.0, https://github.com/hzi-bifo/Haploflow
  - PEHaplo [6]: 1.0, https://github.com/chjiao/PEHaplo
  - VStrains: v0.0.1, https://github.com/MetaGenTools/VStrains

##### Software commands

*Read trimming and adapter removal for real datasets*
fastp [7]: fastp -i 1.fastq -o 1.fp.fastq -I 2.fastq -O 2.fp.fastq -e 36

*Read alignment and preprocess for 2-SARS-COV-2 real datasets*

1. Individually assemble ground-truth reference sequences using SPAdes [4]
   - python spades.py -1 SRR18009684_1.fastq -2 SRR18009684_2.fastq --careful -t 32 -o <output_dir>
   - python spades.py -1 SRR18009686_1.fastq -2 SRR18009686_2.fastq --careful -t 32 -o <output_dir>
2. Generate sam file using BWA [12]
   - bwa index contigs.fasta
   - bwa mem contigs.fasta 1.fq 2.fq > <prefix>.sam
3. Generate sorted bam file using SAMtools [14]
   - samtools view -S -b <prefix>.sam > <prefix>.bam
   - samtools sort <prefix>.bam > <prefix>.sorted.bam
   - samtools index <prefix>.sorted.bam
4. Filter reads using SAMtools [14]
   - samtools view -f 0x2 <prefix>.sorted.bam -o <prefix>.sorted.fil.bam -h <best_contig_name>
   - samtools sort -n <prefix>.sorted.fil.bam -o <prefix>.2sorted.fil.bam
5. Convert back to pair-end fastq format using SAMtools [14]

  – samtools fastq -@ 8 <prefix>.2sorted.fil.bam -1 <output_name>.forward.fastq -2 <output_name>.reverse.fastq -N

*De novo assembly tools*

- SPAdes-series [1,4,5,15,17]
  - python spades.py -1 forward.fastq -2 reverse.fastq --careful -t 32 -o <output_dir>
  - python spades.py -1 forward.fastq -2 reverse.fastq --meta -t 32 -o <output_dir>
  - python spades.py -1 forward.fastq -2 reverse.fastq --metaviral -t 32 -o <output_dir>
  - python spades.py -1 forward.fastq -2 reverse.fastq --rnaviral -t 32 -o <output_dir>
  - python coronaspades.py -1 forward.fastq -2 reverse.fastq -t 32 -o <output_dir>
- SAVAGE [2]
  - haploconduct savage -p1 forward.fastq -p2 reverse.fastq --revcomp --split 30 -t 32
- VG-Flow [3]
  - python build_graph_msga.py -f forward.fastq -r reverse.fastq -c contigs_stage_c.fasta -t 32 -vg vg
  - python vg-flow.py -m 100 -c 200 --greedy_mode=all  node_abundance.txt contig_graph.final.gfa
- PEHaplo [6]
  - For 20,000x coverage datasets: python pehaplo.py -f1 <fwd.fasta> -f2 <rev.fasta> -l 210 -l1 220 -r 250 -F 450 -n 3 -correct yes -t 32
  - For 4,000x coverage datasets: python pehaplo.py -f1 <fwd.fasta> -f2 <rev.fasta> -l 50 -r 70 -n 3 -correct yes -t 32
- viaDBG [8]
  - For datasets with read length=2×250bp: ./bin/viaDBG -p <paired_end_dir> -o <output> -u <unitig> -k 127 -c dsk -n -t 16 --postprocess
  - For datasets with read length=2×75bp: ./bin/viaDBG -p <paired_end_dir> -o <output> -u <unitig> -k 55 -c dsk -n -t 16 --postprocess
- Haploflow [9]
  - haploflow --read-file <combined_single_reads.fastq> --out <output_dir>
- VStrains
  - python VStrains.py -a spades -g assembly_graph_after_simplification.gfa -p contigs.paths -o <output_dir> -fwd forward.fastq -rve reverse.fastq

*Reference-based assembly tools*
PredictHaplo [18]: PredictHaplo-Paired config.txt

*Assembly evaluation*
MetaQUAST [16]: python2 metaquast.py --unique-mapping --report-all-metrics -m 500 -t 8 <f1.fasta> <f2.fasta> ... <fm.fasta> -o <output_dir> -R <ref1.fasta>,<ref2.fasta>,...,<refn.fasta>

**Recorded exceptions in experiments**

1. SAVAGE failed on 2-SARS-COV-2 real dataset, tested with "–split 5/6/7" followed by the instruction (quote from its GitHub page): 500 < read_coverage/patch_num < 1000.
2. PEHaplo failed to complete the final error correction on the 6-Poliovirus simulated datasets.
3. VG-Flow+SPAdes crashed on building variation graph on several datasets when using vg-toolkit (*ERROR: Signal 6 occurred.*), or caught by assertion error traced to line 663 in *vg-flow/scripts/vg-flow.py*.
4. viaDBG incorrectly hard-codes relative path (and typos) in the program, traced to line 6 in *viaDBG/Utils/script/dsk_script_old* and line 347 in *viaDBG/Src/DBG/DBG.h*.
